# Thermal Dynamics of Ectotherm Behavioral Strategies: Insights from Agent-Based Simulations

**DOI:** 10.1101/2025.04.30.651512

**Authors:** Michael J. Remington, Rulon W. Clark, Ryan J. Hanscom, Timothy E. Higham, Jeet Sukumaran

## Abstract

Terrestrial ectotherms use dynamic behavioral thermoregulation strategies that typically rely on finding thermal refugia to avoid unfavorable conditions. Traditional analytic approaches rely on analysis of thermal indices to describe thermal conditions organisms may or may not choose, though computational simulations could potentially be used to analyze thermoregulatory behavior explicitly. Here, we leverage a novel simulation framework that integrates data from operative thermal models deployed in microhabitats available to ectotherms to showcase the trade-offs between two basic thermoregulatory strategies: maintaining a thermal optimum point or attempting to keep their body temperature within a preferred thermal range. We assess the output from simulations of these behavioral strategies to understand their influence on body temperature regulation and occupancy of thermal refugia, and then compare our findings from an empirical case study. Results from our analysis suggest that these two general strategies do not impact thermoregulation as strongly as previously assumed. We also found that occupancy of thermal refugia appeared to be driven by environmental conditions rather than the thermoregulation strategy. Although our case study showed promise in providing insights into predicting use of thermal refugia, understanding microhabitat occupancy from thermal data alone may require finer scale temporal data, or the integration of other factors such as foraging dynamics or predation avoidance.

**Highlights:**

- Agent-Based simulation framework which explicitly models ectotherm behavioral thermoregulation.
- Analysis shows minimal differences between varying thermoregulation strategies.
- Simulations identify cathermal activity patterns regardless of behavioral strategy.
- Results show promise of empirical applications.

## 1. Introduction

Behavioral thermoregulation is critical for ectothermic organisms, which depend on external heat sources and are highly susceptible to environmental temperature fluctuations. Microhabitats in the landscape may vary in thermal conditions across both space and time, thus ectotherms employ dynamic behavioral strategies to find thermal refugia that best maintain their body temperature. For example, terrestrial ectotherms that are semi-fossorial, make or use underground burrows that serve as both thermal and physical refugia that protect them from unfavorable biotic and abiotic conditions (Kinlaw (1999)). Since body temperature regulates enzymatic processes (Kern et al. (2015)) and muscle function (Bennett (1984)), behavioral thermoregulation is essential for optimizing performance in activities like foraging and predator avoidance.

Although some ectotherms exhibit little or no behavioral thermoregulation, most species maintain body temperatures within a preferred thermal range, and some may regulate body temperature more precisely near a thermal optimum (Brown and Weatherhead (2000), Hertz et al. (1993), Piasečná et al. (2015)). The spectrum of behavioral strategies involved in thermoregulation has numerous consequences on evolutionary fitness and ecological interactions. Because hiding in refugia almost always involves a trade-off with other fitness-critical activities (e.g., finding food or mates), the movement of individuals back and forth between refugia and more open microhabitats is a central aspect of the daily behavioral cycle of terrestrial ectotherms.

Research on ectotherm thermal ecology has primarily relied on mathematical models including dynamic energy budget theory or other niche-based model. However, these approaches typically focus on population-level trends, potentially overlooking behavioral variability at the individual level (Sears and Angilletta (2015)), furthermore, some mathematical models have been criticized for their lack of capacity to capture fine-scale spatio-temporal temperature conditions (Kearney and Porter (2009)). Along with this critique, an ectotherm’s body temperature condition is not only dependent on its current microhabitat but also lagging microhabitat states the individual sampled Ha et al. (2011), thus aggregating thermal conditions across microhabitats may fail to capture dynamics related to behavioral thermoregulation.

While traditional field-based methods such as camera traps and field surveys provide valuable ecological insights, they are labor-intensive and limited in their ability to capture large volumes of temporally granular behavioral data (Rowcliffe et al. (2008), Young et al. (2018)). Another prominent approach in thermal ecology involves characterizing thermoregulatory behavior using operative temperature models (OTMs), which continuously sample temperatures across available microhabitats. These data are then used to calculate heuristic thermal indices such as thermal accuracy, thermal quality, and thermal effectiveness (Taylor et al. (2021)).

OTM-based methods rely on operative temperature (Te), measured using physical OTMs, to infer thermal environments experienced by ectotherms. Understanding how ectotherms sample various microhabitats with different Te values can be approached through mathematical models (Dzialowski (2005)) or summarized in reference to specific microhabitats (Crowell et al. (2021)). However, rather than aggregating across microhabitats, reducing multidimensional Te vectors through simulation into a single dynamic trajectory could provide deeper insights into individual thermal refuge selection. For example, the thermal conditions of a basking lizard can be represented as Te*_r_ock* (rock surface temperature) versus Te*_shade_*(shade temperature). A simulation model could simulate the lizard’s chosen environment at any given time point; resulting in a vector of Te*_lizard_*, which reflects the thermal conditions it experiences based on its behavioral choices.

Here, we explore the utility of a novel agent-based thermoregulatory simulation framework designed to explicitly model the behavioral strategies ectotherms use to navigate thermal variability in their habitat. Our analysis proceeds along two lines: first, we conceptually quantify the thermal trade-offs that arise from different thermoregulatory strategies; second, we demonstrate the utility of our approach through a case study, applying the framework to empirical data and showing how it can supplement current OTM based studies. By doing so, we demonstrate our tool has the potential not only to predict individual body temperature patterns and microhabitat occupancy, but also to offer thermal ecologists a more comprehensive means of understanding and quantifying behavioral strategies. Ultimately, this framework can help classify which strategies ectotherms adopt under varying environmental conditions and potential estimate the rates at which they occupy thermal refugia, thereby improving our insight into daily activity patterns and habitat thermal quality.

## 2. Methods

Our simulation software is an agent-based simulation model (entitled ThermaNewt, as it relies on Newton’s Law of Cooling) that simulates body temperature changes and microhabitat occupancy using time series data from temperature loggers placed in key microhabitats available to ectotherms. To see the full analysis pipeline, refer to Figure 1. The model simulations and post simulation analysis were implemented in Python 3.8.5 (Van Rossum and Drake (2009)).

**Figure 1:**
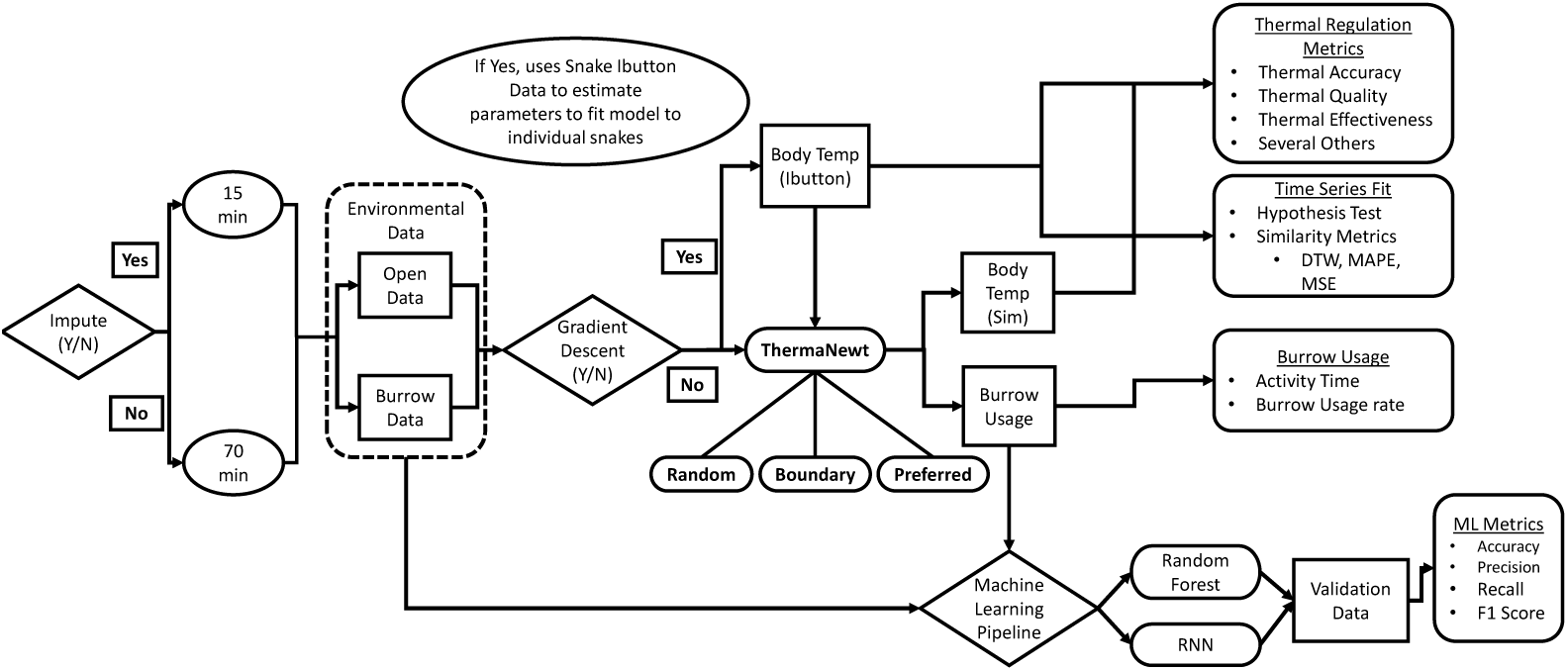
Workflow of the simulation analysis conducted. ThermaNewt, is the name for the simulation model based Newton’s law of cooling. Random, Boundary, and Preferred represent our three thermoregulation sub-models (Figure 2).

### 2.1. Empirical Data

For an empirical application of our approach, we used a field study of three populations of Prairie Rattlesnakes (*Crotalus Viridis*) located at North American sites across a latitudinal cline (Canada site: South Eastern Alberta Canada in the Palliser Triangle, Nebraska site: Northern Nebraska near the Niobrara River in the Central Great Plains ecoregion, Texas Site: Near the city of Marathon in the Chihuahuan Desert). Individual snakes at these sites were surgically implanted with radio transmitters (Wildlife Materials, SOPI-2380-MVS) and Thermochrom iButtons DS1922L-F5, accuracy = ±0.5°C, Maxim Integrated Products Inc., Rio Robles, San Jose, CA), which recorded field-active body temperatures (Tb) every 70 minutes. Snakes were released 1-2 days following surgery at the location of capture. Snakes were recaptured after one year of monitoring and the iButton and radio transmitter were surgically removed. Our model incorporated 10 snakes from the Canada site, 2 from the Nebraska site, and 9 from the Texas site. At each site we also placed three replicate physical Operative Thermal Models (OTMs) in each of four different microhabitats typically used by this species (inside burrow, at entrance of burrow, under vegetation, and on open ground); however, to generate this initial case study, we simplified this array of microhabitats to the two extremes (i.e., the two microhabitat most likely to differ in temperature): in burrow, and in open. OTMs consisted of hollow copper tubes that replicate thermal qualities of snake bodies (Alujević et al. (2024)).

We were also able to intermittently validate microhabitat occupancy of snakes from fixed videography and direct observation via individuals with telemeters used in the field to monitor snake behavior. This provides sparse, semi-continuous time series observations of whether the focal snake was in the burrow or out in the open. This data was used to evaluate the accuracy of our simulated microhabitat occupancy classification output from our model.

### 2.2. Simulation Model Description

The simulation model is based on a discrete, time-dependent version of Newton’s Law of Cooling (Equation (1)), which describes the change in body temperature over time due to heat exchange with the environment. The framework uses thermal time series data from temperature loggers that are exposed to the direct sun or placed in thermal refugia that could potentially be occupied by a focal species, and then simulates body temperature changes and occupancy of thermal refugia. During each cycle, the internal body temperature of the agent is calculated via Newton’s law of cooling equation with respect to time.

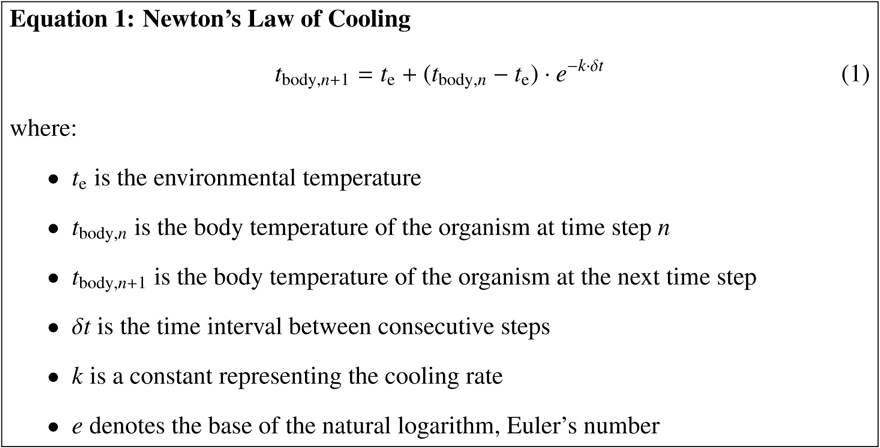

Newton’s law of cooling has a rich history in the study of organismal thermal ecology (Scholander et al. (1950), Stevenson (1985), Dreisig (1985)). Because *T_e_* often differs between potential microhabitats, we developed behavioral response functions as sub-models that relate the agent’s internal body temperature to the probability of the organism switching microhabitat occupancy (open or refuge) to infer which microhabitat temperature the organism occupies at any given time step.

We developed three basic thermoregulatory sub-models: *random*, *boundary*, and *preferred* (Figure 2), and a fourth sub-model as a variant of the *preferred* sub-model (*preferred_est_*). The *random* sub-model serves as a null model where agents switch microhabitat occupancy randomly, regardless of their internal temperature. In the *boundary* sub-model, agents change occupancy when their internal temperature falls outside a defined thermal preference range. Finally, in the *preferred* sub-model, agents try to maintain their body temperature near a defined thermal optimum, with the probability of switching states increasing as their temperature deviates from this optimum. The preferred*_est_* is a variant of the preferred sub-model which models intraspecific differences in thermal preferences by estimating thermal preference variables from empirical data (see Parameter Estimation with Gradient Descent section for more details)

**Figure 2:**
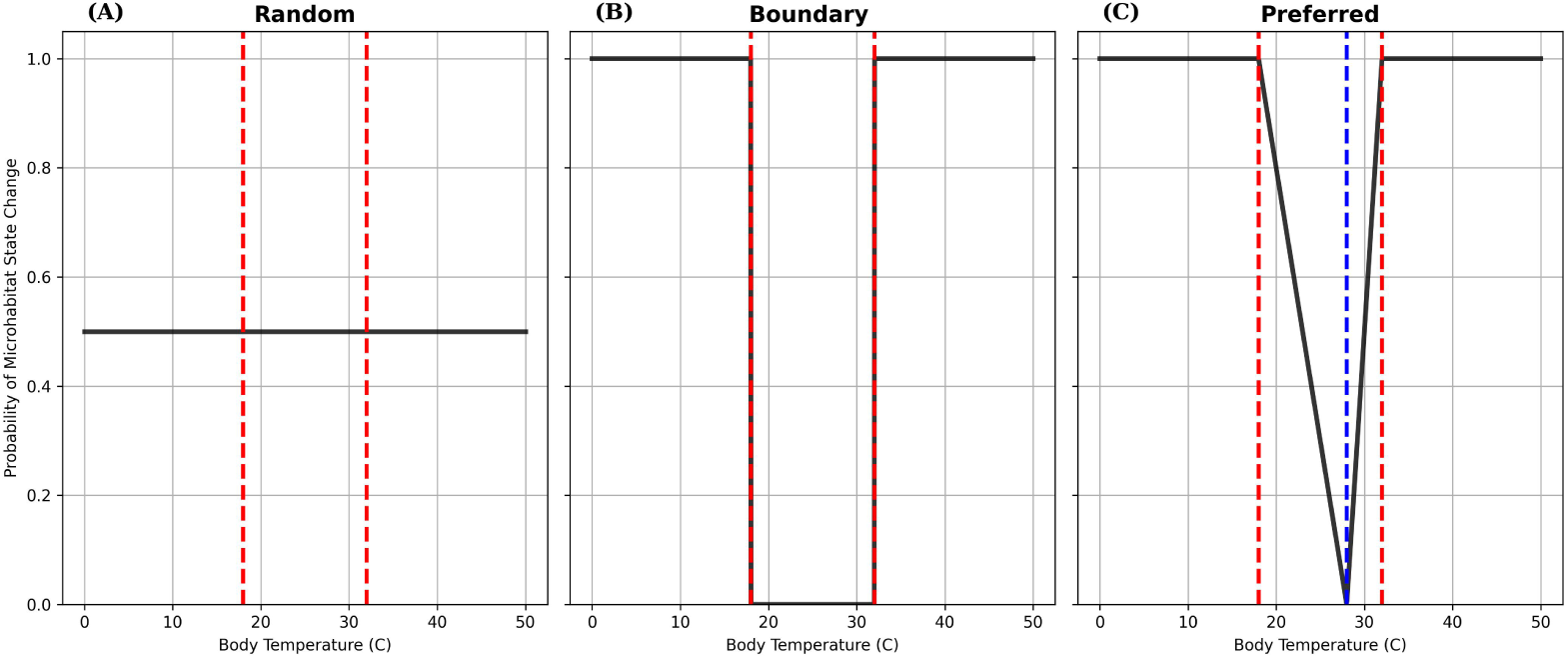
Behavioral response functions for our three thermoregulatory sub-models. The x axis is the organism’s internal body temperature and the y axis is the probability of an agent switching to the optimal thermal refugia. Red lines represent the thermal preference range we set for the snakes. A) Represents the random model, where our modeled organism flips states at random regardless of internal body temperature. B) The boundary model where the organism changes state if it is no longer in the thermal preference range. C) The preferred model represents an organism trying to maintain its body temperature along a thermal optima (shown in blue).

Although empirical T*_set_* measurements (i.e., the range of body temperatures within which an ectotherm exhibits optimal physiological performance) for the focal populations of our case study *Crotalus viridis* were not available from an experimental design, but most observations indicate that their summer body temperatures range between 20–30°C (Redder (1994)). Experimental data for a closely related species (*C. oreganus*) indicate that minimum T*_set_* values can vary by population but typically lie between 11–16°C, while maximum T*_set_* values range from 36–38°C (Crowell et al. (2021)). Based on this information and under the assumption an ectotherms thermal preference range is a subset of it’s T*_set_*, we assumed a conservative minimum thermal preference of 18°C and a maximum thermal preference of 32°C for our model. Parameters for the random, boundary, and preferred sub-models are provided in Table 1. Estimated thermal preference parameters for the preferred*_est_* sub-model are listed in Table 2.

**Table 1:** Parameter values for each of sub-model when parameters are not estimated via the gradient descent algorithm. For the preferred*_est_* case, parameters for thermal minimum, thermal maximum, and thermal optima vary for each rattlesnake (see supplementary materials).

| Sub-Model | $t_{\text{initial}}$ | k | $T_{\text{pref.min}}$ | $T_{\text{pref.max}}$ | $T_{\text{opt}}$ |
| --- | --- | --- | --- | --- | --- |
| Random | 25 | 0.01 | - | - | - |
| Boundary | 25 | 0.01 | 18 | 32 | - |
| Preferred | 25 | 0.01 | 18 | 32 | 28 |
| Preferred<sub>est</sub> | 25 | 0.01 | - | - | - |

**Table 2:**
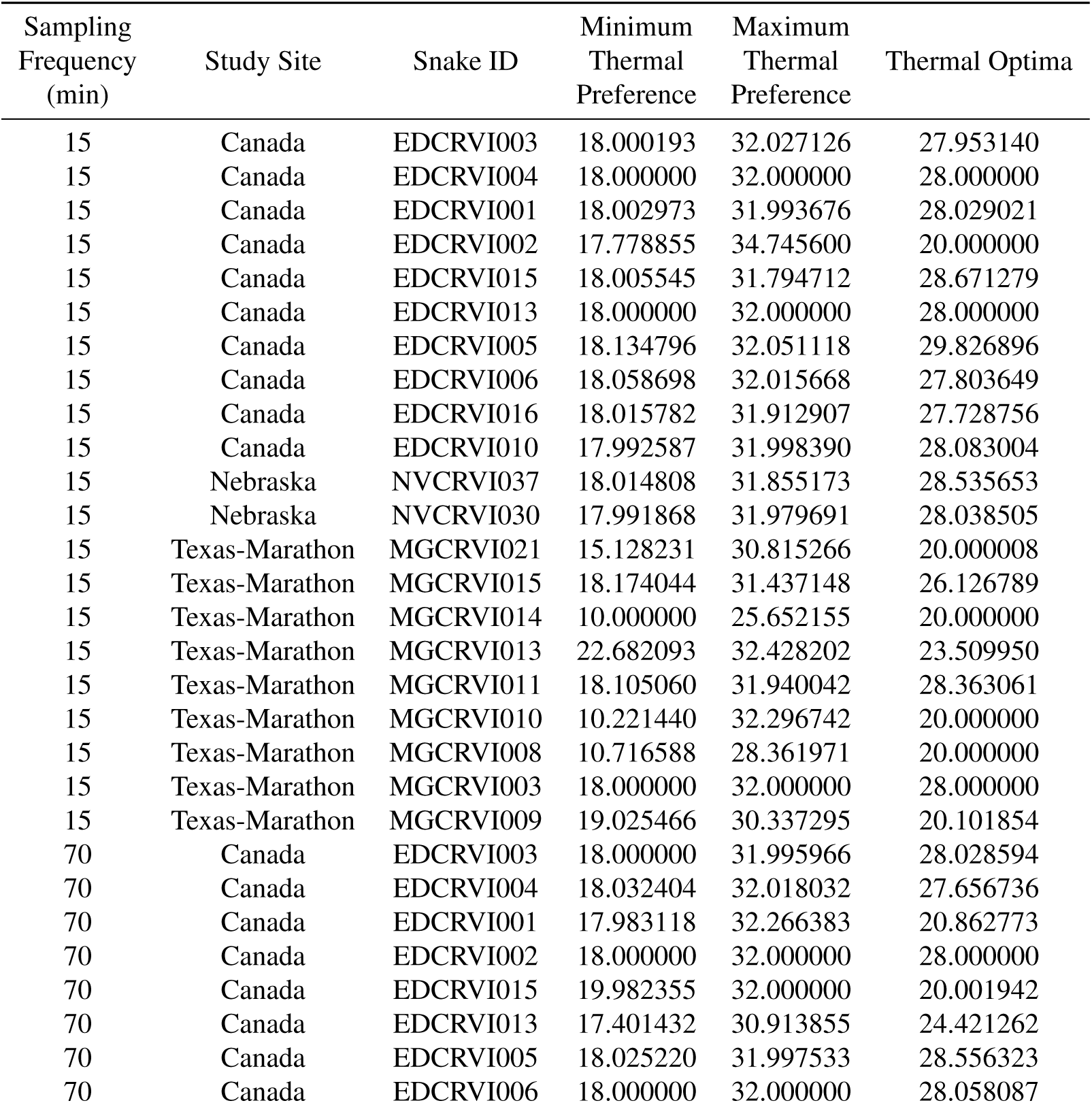
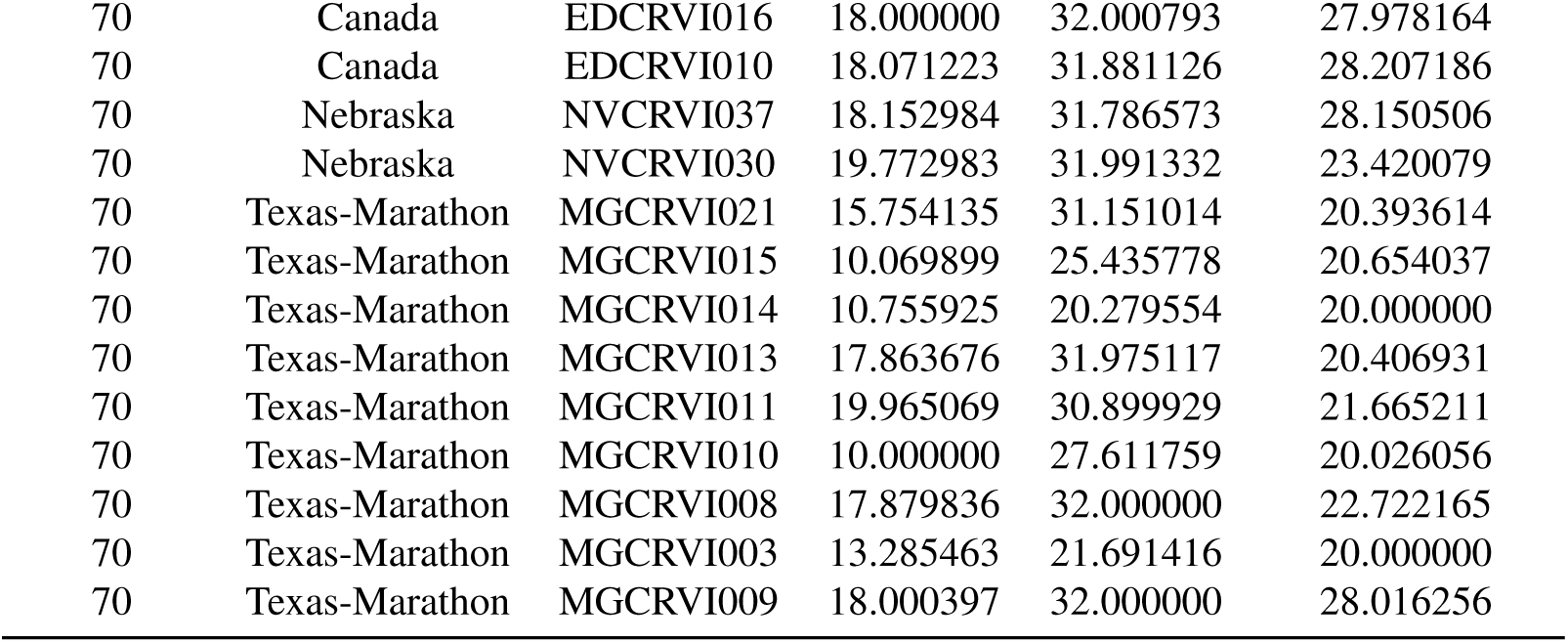
Estimated thermal preference values used in the preferred*_est_* sub-model. Data with a sampling frequency of 70 minutes was not imputed, whereas data with a sampling frequency of 15 minutes was imputed.

| Sampling Frequency (min) | Study Site | Snake ID | Minimum Thermal Preference | Maximum Thermal Preference | Thermal Optima |
| --- | --- | --- | --- | --- | --- |
| 15 | Canada | EDCRVI003 | 18.000193 | 32.027126 | 27.953140 |
| 15 | Canada | EDCRVI004 | 18.000000 | 32.000000 | 28.000000 |
| 15 | Canada | EDCRVI001 | 18.002973 | 31.993676 | 28.029021 |
| 15 | Canada | EDCRVI002 | 17.778855 | 34.745600 | 20.000000 |
| 15 | Canada | EDCRVI015 | 18.005545 | 31.794712 | 28.671279 |
| 15 | Canada | EDCRVI013 | 18.000000 | 32.000000 | 28.000000 |
| 15 | Canada | EDCRVI005 | 18.134796 | 32.051118 | 29.826896 |
| 15 | Canada | EDCRVI006 | 18.058698 | 32.015668 | 27.803649 |
| 15 | Canada | EDCRVI016 | 18.015782 | 31.912907 | 27.728756 |
| 15 | Canada | EDCRVI010 | 17.992587 | 31.998390 | 28.083004 |
| 15 | Nebraska | NVCRVI037 | 18.014808 | 31.855173 | 28.535653 |
| 15 | Nebraska | NVCRVI030 | 17.991868 | 31.979691 | 28.038505 |
| 15 | Texas-Marathon | MGCRVI021 | 15.128231 | 30.815266 | 20.000008 |
| 15 | Texas-Marathon | MGCRVI015 | 18.174044 | 31.437148 | 26.126789 |
| 15 | Texas-Marathon | MGCRVI014 | 10.000000 | 25.652155 | 20.000000 |
| 15 | Texas-Marathon | MGCRVI013 | 22.682093 | 32.428202 | 23.509950 |
| 15 | Texas-Marathon | MGCRVI011 | 18.105060 | 31.940042 | 28.363061 |
| 15 | Texas-Marathon | MGCRVI010 | 10.221440 | 32.296742 | 20.000000 |
| 15 | Texas-Marathon | MGCRVI008 | 10.716588 | 28.361971 | 20.000000 |
| 15 | Texas-Marathon | MGCRVI003 | 18.000000 | 32.000000 | 28.000000 |
| 15 | Texas-Marathon | MGCRVI009 | 19.025466 | 30.337295 | 20.101854 |
| 70 | Canada | EDCRVI003 | 18.000000 | 31.995966 | 28.028594 |
| 70 | Canada | EDCRVI004 | 18.032404 | 32.018032 | 27.656736 |
| 70 | Canada | EDCRVI001 | 17.983118 | 32.266383 | 20.862773 |
| 70 | Canada | EDCRVI002 | 18.000000 | 32.000000 | 28.000000 |
| 70 | Canada | EDCRVI015 | 19.982355 | 32.000000 | 20.001942 |
| 70 | Canada | EDCRVI013 | 17.401432 | 30.913855 | 24.421262 |
| 70 | Canada | EDCRVI005 | 18.025220 | 31.997533 | 28.556323 |
| 70 | Canada | EDCRVI006 | 18.000000 | 32.000000 | 28.058087 |
| 70 | Canada | EDCRVI016 | 18.000000 | 32.000793 | 27.978164 |
| 70 | Canada | EDCRVI010 | 18.071223 | 31.881126 | 28.207186 |
| 70 | Nebraska | NVCRVI037 | 18.152984 | 31.786573 | 28.150506 |
| 70 | Nebraska | NVCRVI030 | 19.772983 | 31.991332 | 23.420079 |
| 70 | Texas-Marathon | MGCRVI021 | 15.754135 | 31.151014 | 20.393614 |
| 70 | Texas-Marathon | MGCRVI015 | 10.069899 | 25.435778 | 20.654037 |
| 70 | Texas-Marathon | MGCRVI014 | 10.755925 | 20.279554 | 20.000000 |
| 70 | Texas-Marathon | MGCRVI013 | 17.863676 | 31.975117 | 20.406931 |
| 70 | Texas-Marathon | MGCRVI011 | 19.965069 | 30.899929 | 21.665211 |
| 70 | Texas-Marathon | MGCRVI010 | 10.000000 | 27.611759 | 20.026056 |
| 70 | Texas-Marathon | MGCRVI008 | 17.879836 | 32.000000 | 22.722165 |
| 70 | Texas-Marathon | MGCRVI003 | 13.285463 | 21.691416 | 20.000000 |
| 70 | Texas-Marathon | MGCRVI009 | 18.000397 | 32.000000 | 28.016256 |

### 2.3. Data Integration and Interpolation

All data analysis and cleaning were conducted using Pandas (The pandas development Team (2024)), a python library for dataframe manipulation. We averaged the microhabitat temperature data across OTMs placed in replicate microhabitats to the nearest hour, and matched it to the corresponding snake body temperature timeline using Pandas’ *merge aso f* function with the forward-filling method. This approach creates a unified timeline for all temperature variables at each site, associating snake body temperatures with climate observations that occurred within a 70-minute window.

Data from cooler months (October through April) when snakes are inactive were excluded from the analysis. Snake body temperatures during overwintering remain nearly constant and are often higher than near-surface conditions because snakes use deeper underground refugia during these months.

### 2.4. Parameter Estimation with Gradient Descent

One assumption of our base analysis is that there is no intraspecific variation in thermal preferences between populations of rattlesnakes. This assumption may not reflect real world conditions as evidence exists that individuals of some species of ectotherms can have variance in thermal preferences (Nati et al. (2021)).

To evaluate the impact of accounting for individual variation, we relaxed this assumption and conducted a separate preferred sub-model (preferred*_est_*). This analysis fits our model to observed rattlesnake body temperature data and estimating the minimum, maximum, and optimal thermal preferences using a gradient descent algorithm.

If this model fits the empirical data significantly better than our other models, it would suggest that individual differences in thermal preference play a greater role in driving thermoregulation dynamics than the specific thermoregulatory strategy itself. Conversely, if the model performs similarly or worse, it would imply that individuals across the population adopt static strategies, with thermal preferences being less influential in thermoregulation.

We used a bounded limited memory BFGS ‘L-BFGS-B’ optimization algorithm with an approximated gradient with a step size of 0.01 to estimate parameters. We found this method outperformed the adam optimization algorithm at recovering simulated parameters. Estimated parameters used in analysis can be seen in Table 2. Note that this sub-model was constructed to demonstrate the potential utility that a simulation approach has for considering individual variation in thermoregulatory strategies, and that future use of this approach would need to incorporate estimates of individual variation in body temperature preferences that were independent from field-collected data (i.e., collected from animals occupying a thermal gradient under controlled conditions).

### 2.5. Thermal Indices

The effectiveness of ectotherm thermoregulation has historically been evaluated through heuristics based on thermal indices. Our simulation software outputs simulated ectotherm body temperature (T*_b_*), and the environmental conditions of the agent as a function of microhabitat occupancy (open or burrow). For each of our thermoregulatory sub-models, we evaluate four thermal indices traditionally used in thermal ecology studies: thermal quality (de), thermal accuracy (db), thermal effectiveness (E), and the thermoregulatory exploitation index (Ex) (Hertz et al. (1993), Taylor et al. (2021)).

Thermal accuracy (db) is a metric that measures the difference between an organism’s body temperature (T*_b_*) and its thermal preference zone. High values of db indicate poor thermal accuracy. Thermal quality (de) is a metric that measures the distance of the thermal operative temperature (T*_e_*) from the thermal preference zone, with high values of de indicating low quality thermal habitat. Thermal effectiveness (E) is one minus the ratio of mean db and mean de. Values of 1 indicate careful and precise thermoregulation, and 0 indicates that the animal is thermoconforming (body temperature always matches the environmental temperature). Values approaching negative 1 or lower indicate periods of where the organism is in poor habitat and are experiencing thermal stress. Finally, the thermal exploitation index (Ex) is the percentage of time spent in the thermal preference zone, with higher values indicating more time spent in the thermal preference zone (Hertz et al. (1993), Piasečná et al. (2015), Taylor et al. (2021))

### 2.6. Time Series Comparison Analysis

One of the primary objectives of our simulation is to evaluate which thermoregulatory sub-model best approximates the body temperatures observed in empirical data. To assess the similarity between empirical and simulated data, we conducted a time series comparison analysis. This involved calculating three key distance metrics for each snake: mean squared error (MSE), mean absolute percentage error (MAPE), and dynamic time warping distance (DTW) (Berndt and Clifford (1994)). MSE quantifies average squared differences between simulated and empirical temperatures, highlighting large deviations. MAPE measures prediction accuracy as a percentage, offering interpretability across scales. DTW captures temporal alignment by comparing time series with potential shifts in timing, making it ideal for assessing sequences with time lags or irregular patterns. These metrics collectively provide a comprehensive evaluation of both prediction accuracy and temporal correspondence.

We assume that the sub-model which minimizes these distance metrics represents the most probable thermoregulatory strategy adopted by free-ranging rattlesnakes. For each distance metric, we applied a mixed linear model to statistically determine whether a behavioral sub-model significantly reduced these metrics compared to a random sub-model. The study site location was included as a fixed effect, and individual snakes were treated as random effects in each model.

### 2.7. Predicting Microhabitat Occupancy with Supervised Machine Learning

We used a supervised machine learning pipeline to assess which sub-model best approximates empirical data and determine if we could accurately predict microhabitat occupancy (burrow usage) using our simulation framework. We performed an exhaustive search of different machine learning parameter combinations to evaluate the accuracy of our method in predicting ectotherms’ occupancy of thermal refugia. If our pipeline consistently achieves high accuracy, it could become a valuable tool for predicting population-level patterns such as daily activity times or phenology.

Two machine learning algorithms were evaluated, a random forest (RF) approach (Breiman (2001)) and a recurrent neural network (RNN) approach (Elman (1990)). We chose these two methods because RNNs are specifically designed for sequential data, making them well-suited for time series forecasting. While RFs are not inherently designed for sequential data, they can still be effective in time series analysis when temporal features are explicitly engineered.

For the RF approach, we used the month, hour, open temperature, and burrow temperature as our explanatory features and used simulated burrow occupancy labels from the thermal model as our response variable for the classification algorithm. Hyper parameters were estimated via a randomized hyper-parameter optimization algorithm called *RandomizedS earchCV* (Pedregosa et al. (2011)). Hyper-parameters are parameters that do not have biological relevance but impact model fit. Data were partitioned into a training and a test set (80/20) rate.

The RNN pipeline parsed our training and test data into sequences. We tested window sizes (3-10) to see which window size was optimal. Empirical behavioral validation data from video analysis was also parsed into sequences to make them directly comparable.

For both models, we evaluate accuracy, precision, recall, and F1 scores against an empirical validation set outside of the training and test sets of the machine learning model, using functions provided by the scikit-learn library (Pedregosa et al. (2011)). We apply both models to all combinations of our simulation pipeline including imputing versus not imputing data, each of the thermoregulatory sub-models, parameters estimated (yes or no).

## 3. Results

### 3.1. Ecological Comparison of Thermoregulatory Strategy

Among our thermoregulatory study groups (random, boundary, preferred, and preferred*_est_*), we found that the boundary thermoregulatory sub-model led to the highest rate of exposure (agents occupying open habitat), while also having more state changes compared to the preferred. Both the preferred and the preferred*_est_* models resulted in lower exposure rates and fewer state changes than the random model (Figure 3A, B).

**Figure 3:**
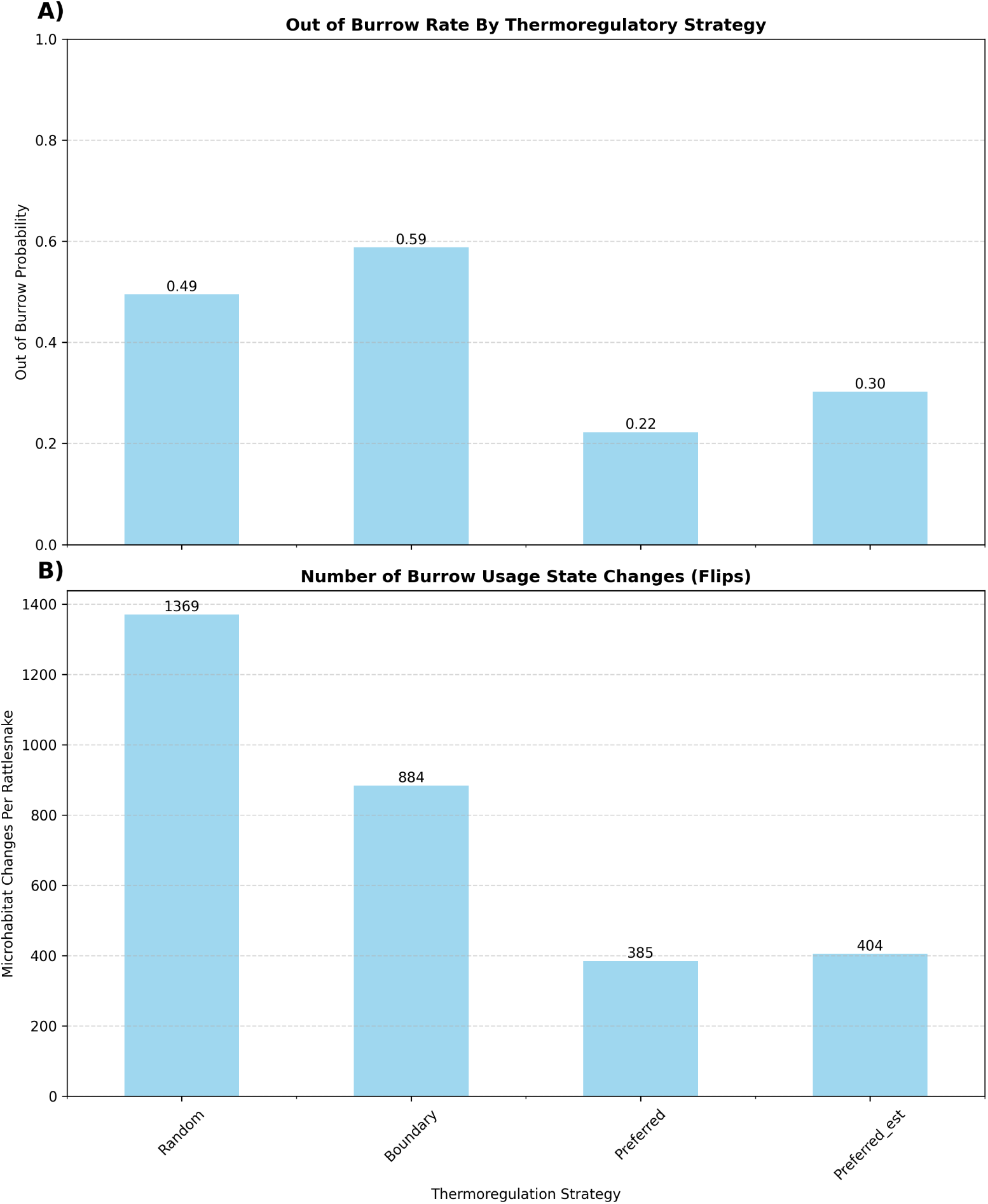
Both graphs showcase microhabitat usage metrics by thermoregulation strategy. Figure A, the y-axis represents the probability of being in the open, where 1 is 100% probability of being in the open and 100% probability of being in the burrow. Figure B is the average count of state flips from being in the open to back to the burrow or vice versa per snake.

Our analysis of 24-hour exposure rates revealed distinct patterns across three agent models. Agents with the boundary, preferred, and preferred*_est_* all demonstrated crepuscular activity patterns, with the probability of being in the open peaking near dawn and dusk. Agents with the boundary model spend more time exposed in the open at all times of a diel cycle compared to the preferred model. Agents encoded with the preferred sub-model show somewhat similar temporal trends, but to a lesser degree, and the preferred*_est_* agents were intermediate between the two (Figure 6).

Thermal index analysis across different times of the day corroborates the idea that simulated agents favor crepuscular activity. During the morning and evening times of the day, thermal quality and thermal accuracy are minimized while thermal exploitation is maximized (Figure 9). These times also correspond to the highest probabilities of being active (Figure 6). This indicated that crepuscular activity is optimal for snakes in reference to maintaining preferred body temperatures; regardless of behavioral strategy. Colder nighttime conditions compel agents to warm up during the morning, but as the day rapidly heats up, they must retreat to thermal refugia to avoid overheating. In the evening, after spending the day sheltered in a burrow, they can become surface active again.

Comparison to empirical data from free-ranging snakes indicates that simulated agents were more likely to occupy open microhabitat than free-ranging snakes. At the Nebraska and Texas study site, the thermal accuracy values of the boundary agents tended to underestimate the thermal accuracy while the preferred sub-model tended to match actual snakes thermal accuracy at those sites (Figure 7 B, C). However, at the Canada site, simulated snake thermal accuracy values more closely matched measures from free-ranging snakes for all sub-models except the random sub-model (Figure 7A).

### 3.2. Simulated Body Temperature Accuracy

Due to the number of snakes and the length of our time series, visually assessing the similarity of all our simulated conditions can be difficult (Figure 4). This subset of data highlights the general finding that the position of the oscillations is accurate, but the overall magnitude of simulated body temperatures do not match actual temperatures.

**Figure 4:**
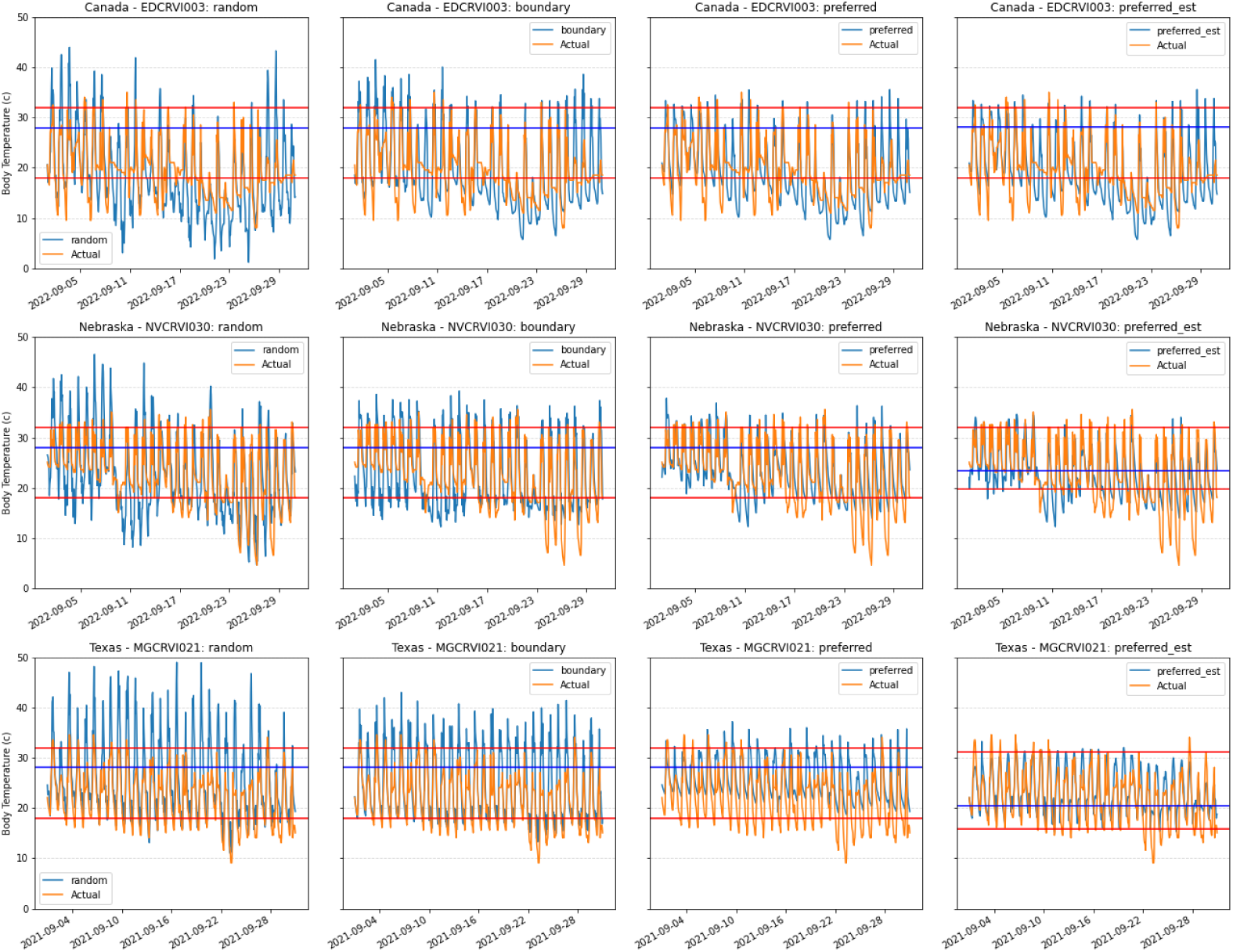
Plot matrix to showcase simulated body temperature of the various thermoregulation strategies (blue) plotted alongside empirical rattlesnake body temperature (yellow) for the month of September. Thermal preference zone marked by red lines. The x axis is the date and the y axis is body temperature. The first row is a snake (EDCRVI003) collected in Canada, the second row is a Nebraska snake (NVCRVI030), and the third is a snake collected in Texas (MGCRVI021). This was with non-imputed data which has a sampling frequency of 70 minutes.

The results of a linear mixed model assessing MSE, MAPE, and DTW between thermoregulatory strategies found the boundary, preferred, and preferred*_est_* all significantly reduced all 3 time series measures compared to the random model, meaning they were more similar to the actual data (Table 3, 4, 5). The preferred sub-model was slightly more effective at minimizing MSE and DTW than the boundary sub-model (Figure 5A, C), while the boundary performed better than the preferred at minimizing MAPE (Figure 5B). The preferred*_est_*performed the best at minimizing time series (Tables 4, 5).

**Figure 5:**
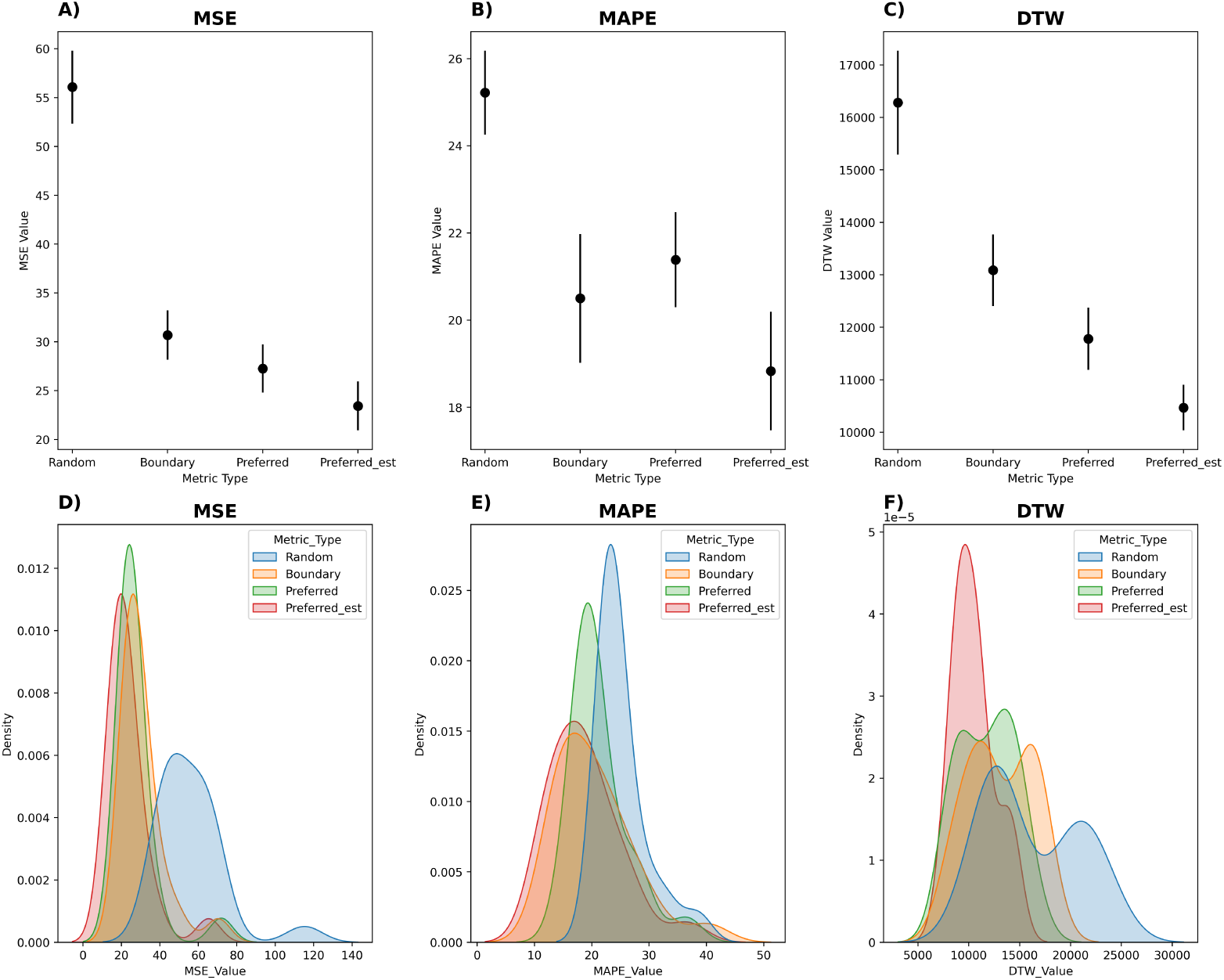
This figure presents a time series distance analysis using Mean Squared Error (MSE), Mean Absolute Percentage Error (MAPE), and Dynamic Time Warping (DTW) for each thermoregulatory strategy. The top row displays the mean and standard error of the time series distance metrics, illustrating the precision and variability of each metric across thermoregulatory sub-models. The bottom row features kernel density distribution plots of the distance values, providing a visual representation of the accuracy distribution for each thermoregulatory strategy. This analysis allows for a quantitative assessment of the effectiveness and consistency of different thermoregulatory sub-models in simulating real-world body temperature dynamics

**Table 3:** Mean least square error (MSE) mixed linear regression results. MSE distances with the thermoregulation strategy as an explanatory variable and study site as a fixed effect. Individual snakes were treated as a random effect. The results of this model suggest that the Boundary, Preferred, and Preferred*_est_* were significantly effective at reducing MSE.

| MSE | Imputed: False |  |  |
| --- | --- | --- | --- |
| Fixed Effects | Coefficients | Std.Err. | P-value |
| Intercept | 53.582 | 3.949 | 0.000*** |
| Strategy: Boundary | -25.383 | 2.197 | 0.000*** |
| Strategy: Preferred | -28.814 | 2.197 | 0.000*** |
| Strategy: Preferred<sub>est</sub> | -32.647 | 2.197 | 0.000*** |
| Study Site: Nebraska | -0.032 | 9.094 | 0.997 |
| Study Site: Texas-Marathon | 5.825 | 5.395 | 0.280 |
| Group Var | 125.174 | 7.358 | - |

**Table 4:** Mean Absolute Percentage Error (MAPE) mixed linear regression results. MAPE distances with the thermoregulation strategy as an explanatory variable and study site as a fixed effect. Individual snakes were treated as a random effect. Based on this model, the simulated body temperatures with parameters estimated (Preferred*_est_*) was the only thermoregulatory sub model that was significantly better at reducing MAPE compared to the random thermoregulatory sub model, which is represented by the intercept group. Overall, study site differences were more significant than the specific thermoregulation strategy sub-model, which could indicate there may be significant differences at the population and individual level in thermoregulation patterns.

| MAPE | Imputed: False |  |  |
| --- | --- | --- | --- |
| Fixed Effects | Coefficients | Std.Err. | P-value |
| Intercept | 28.166 | 1.572 | 0.000*** |
| Strategy: Boundary | -4.718 | 0.625 | 0.000*** |
| Strategy: Preferred | -3.833 | 0.625 | 0.000*** |
| Strategy: Preferred<sub>est</sub> | -6.389 | 0.625 | 0.000*** |
| Study Site: Nebraska | -5.433 | 3.734 | 0.146 |
| Study Site: Texas-Marathon | -5.668 | 2.215 | 0.010** |
| Group Var | 22.214 | 4.363 | - |

**Table 5:** Mixed linear regression results correlating Dynamic Time Warping (DTW) distances with the thermoregulation strategy as an explanatory variable and study site as a fixed effect. Individual snakes were treated as a random effect. Based on this model, the simulated body temperatures with parameters estimated (Preferred*_est_*) was the only thermoregulatory sub model that was significantly better at reducing DTW compared to the random thermoregulatory sub model, which is represented by the intercept group. Overall, study site differences were more significant than the specific thermoregulation strategy, which could indicate there may be significant differences at the population and individual level in thermoregulation patterns.

| DTW | Imputed: False |  |  |
| --- | --- | --- | --- |
| Fixed Effects | Coefficients | Std.Err. | P-value |
| Intercept | 14193.944 | 495.077 | 0.000*** |
| Strategy: Boundary | -3192.678 | 582.384 | 0.000*** |
| Strategy: Preferred | -4499.912 | 582.384 | 0.000*** |
| Strategy: Preferred<sub>est</sub> | -5812.469 | 582.384 | 0.000*** |
| Study Site: Nebraska | -581.110 | 841.114 | 0.490 |
| Study Site: Texas-Marathon | 4998.208 | 498.924 | 0.000*** |
| Group Var | 288795.758 | 237.465 | - |

**Figure 6:**
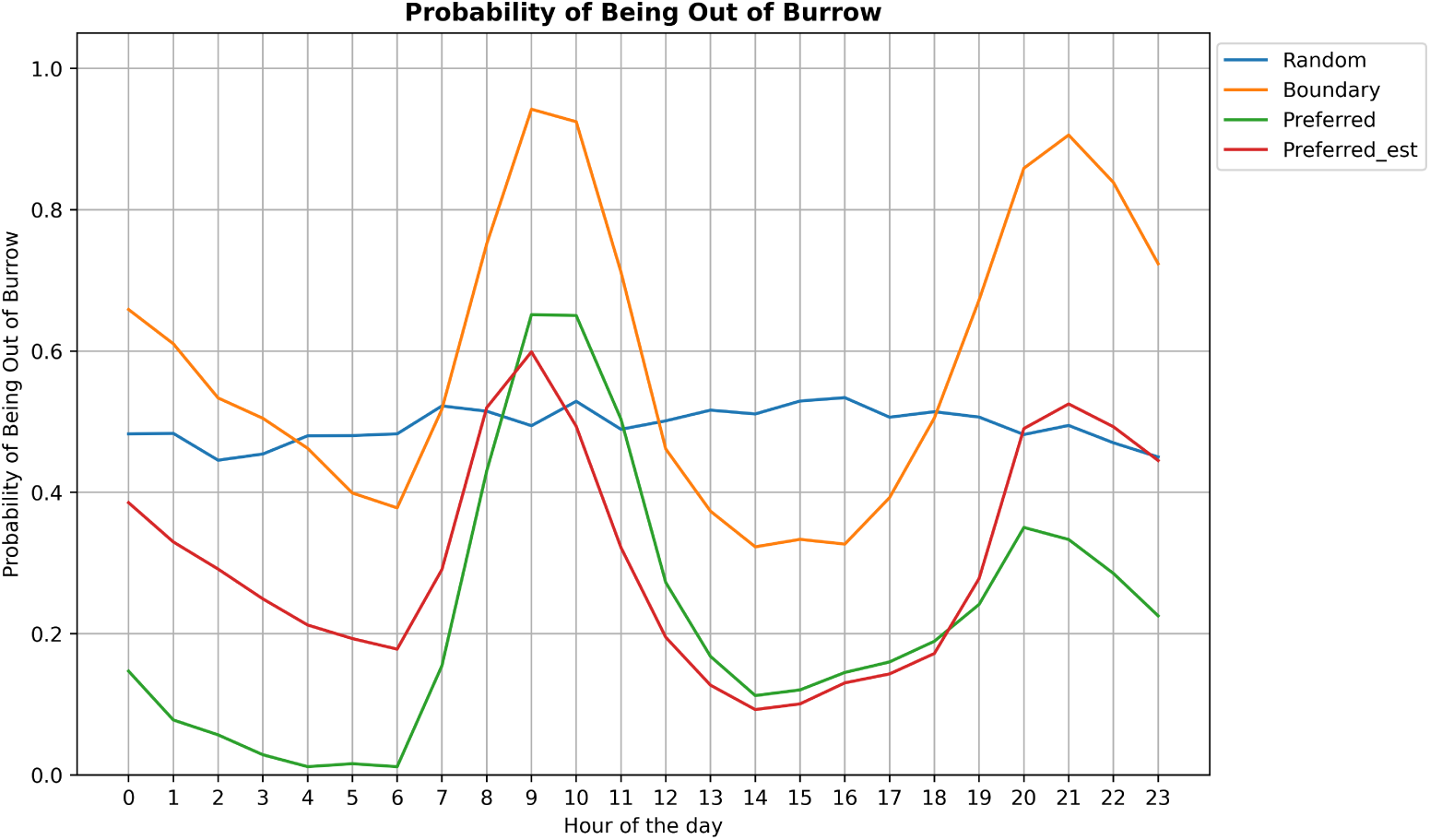
Probability of being in the open by hour. The y-axis represents the probability of being out of the burrow, where 1 is 100% probability of being in the open and 0 is 100% probability of being in the burrow.

**Figure 7:**
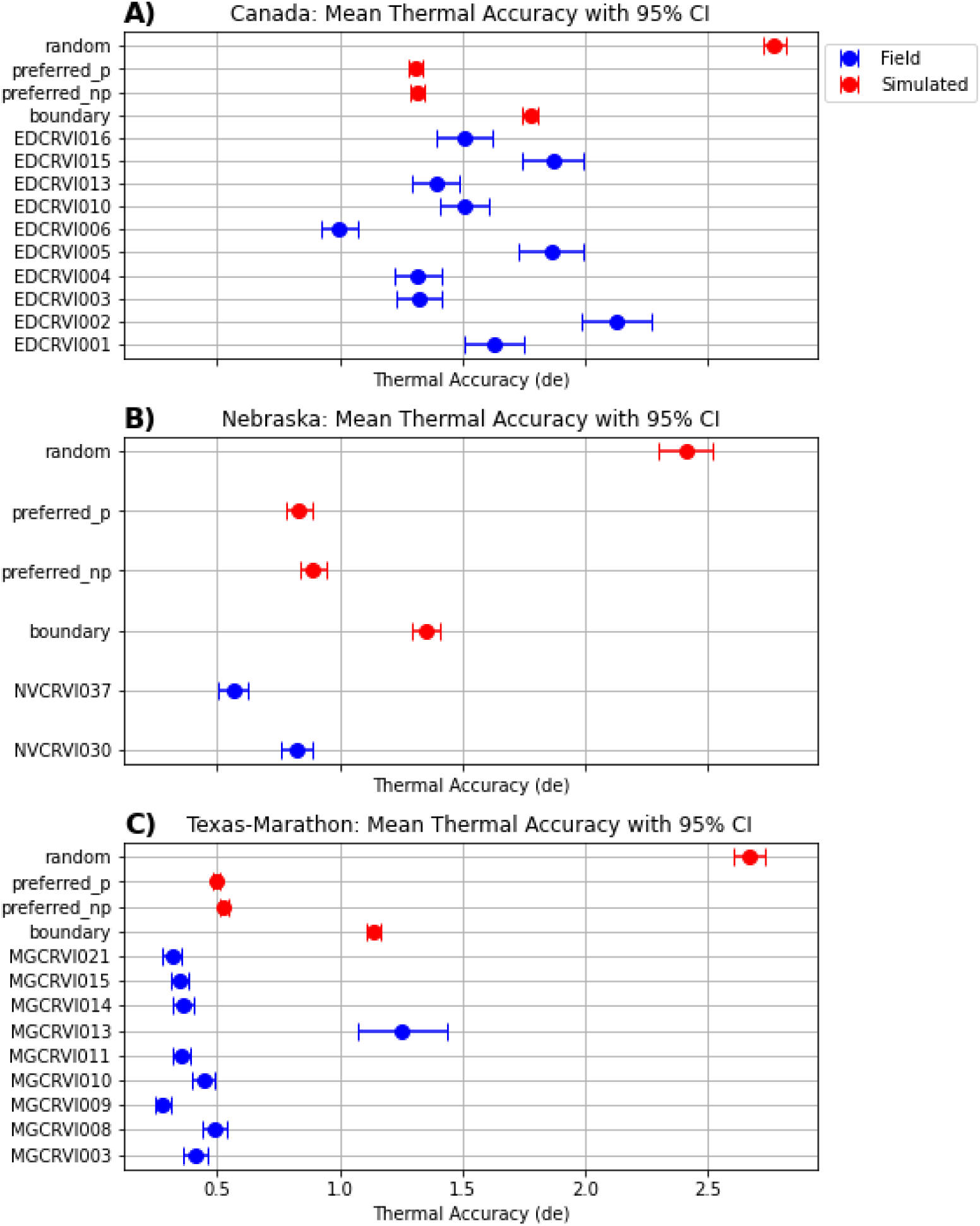
Thermal accuracy for snakes caught in the field, with simulated agents included. Thermal accuracy (db) is a metric that measures the difference between an organism’s body temperature (T*_b_*) from its thermal preference zone. High values of thermal accuracy indicate poor thermal accuracy (Hertz et al. (1993), Piasečná et al. (2015), Taylor et al. (2021)).

### 3.3. Classifying Microhabitat Occupancy Accuracy

From an exhaustive search of different pipeline combinations to evaluate the accuracy of our method in predicting ectotherms’ occupancy of thermal refugia, our most accurate simulation used the preferred thermoregulatory sub-model for the Texas-Marathon study site and yielded an accuracy of 0.717 (Table 6).

**Table 6:** Results of the top 10 most accurate machine learning models predicting microhabitat occupancy of all models ran. Populations were subdivided by the study site and custom models were built from these sub populations marked by the column ‘Study Site’. The first six columns are the model specifications, columns 7-10 showcase model performance, and columns 11-15 are the results of our confusion matrix for each model and AUC which indicates model validity. Metrics such as accuracy, precision, recall, and F1 score are presented alongside additional performance metrics but accuracy is how we sorted the data set to evaluate our top models.

| Top 10 (Accuracy): All Models |  |  |  |  |  |  |  |  |  |  |  |  |  |  |
| --- | --- | --- | --- | --- | --- | --- | --- | --- | --- | --- | --- | --- | --- | --- |
| Study Site | Sub Model | Parameters Estimated? | Sampling Granularity (Min) | ML Method | Window Size | Accuracy | Precision | Recall | F1 Score | TP | FP | TN | FN | AUC |
| Texas-Marathon | preference | Yes | 70 | rf | NaN | 0.716667 | 0.547619 | 0.605263 | 0.575000 | 23 | 19 | 63 | 15 | 0.723363 |
| Texas-Marathon | preference | No | 70 | rf | NaN | 0.691667 | 0.545455 | 0.157895 | 0.244898 | 6 | 5 | 77 | 32 | 0.548460 |
| Texas-Marathon | preference | No | 15 | rf | NaN | 0.690217 | 0.457143 | 0.095238 | 0.157635 | 16 | 19 | 365 | 152 | 0.522732 |
| Texas-Marathon | random | Yes | 15 | rnn | 6.0 | 0.688645 | 0.000000 | 0.000000 | 0.000000 | 0 | 2 | 376 | 168 | 0.431957 |
| Texas-Marathon | random | Yes | 70 | rnn | 8.0 | 0.687500 | 0.560000 | 0.368421 | 0.444444 | 14 | 11 | 63 | 24 | 0.692390 |
| Texas-Marathon | random | No | 70 | rnn | 4.0 | 0.672414 | 0.000000 | 0.000000 | 0.000000 | 0 | 0 | 78 | 38 | 0.688428 |
| Texas-Marathon | random | Yes | 70 | rnn | 9.0 | 0.666667 | 0.509434 | 0.710526 | 0.593407 | 27 | 26 | 47 | 11 | 0.697549 |
| Texas-Marathon | random | Yes | 70 | rnn | 5.0 | 0.660870 | 0.489796 | 0.631579 | 0.551724 | 24 | 25 | 52 | 14 | 0.732057 |
| Texas-Marathon | random | No | 70 | rnn | 8.0 | 0.660714 | 0.500000 | 0.657895 | 0.568182 | 25 | 25 | 49 | 13 | 0.669630 |
| Texas-Marathon | random | Yes | 70 | rnn | 7.0 | 0.654867 | 0.490196 | 0.657895 | 0.561798 | 25 | 26 | 49 | 13 | 0.652281 |

However, when we analyze the distribution of model accuracy grouped by training data population used (Figure 10A), the Texas population has an bimodal distribution while all other accuracy result distributions for the study site and the thermoregulatory sub-models were primarily normally distributed around 0.5 accuracy (i.e., similar to random) (Figure 10A,B). When we assessed the top 10 models using the total population of individuals sampled our best model yielded an accuracy of 0.61 using the random thermoregulatory sub-model (Table 7).

**Figure 8:**
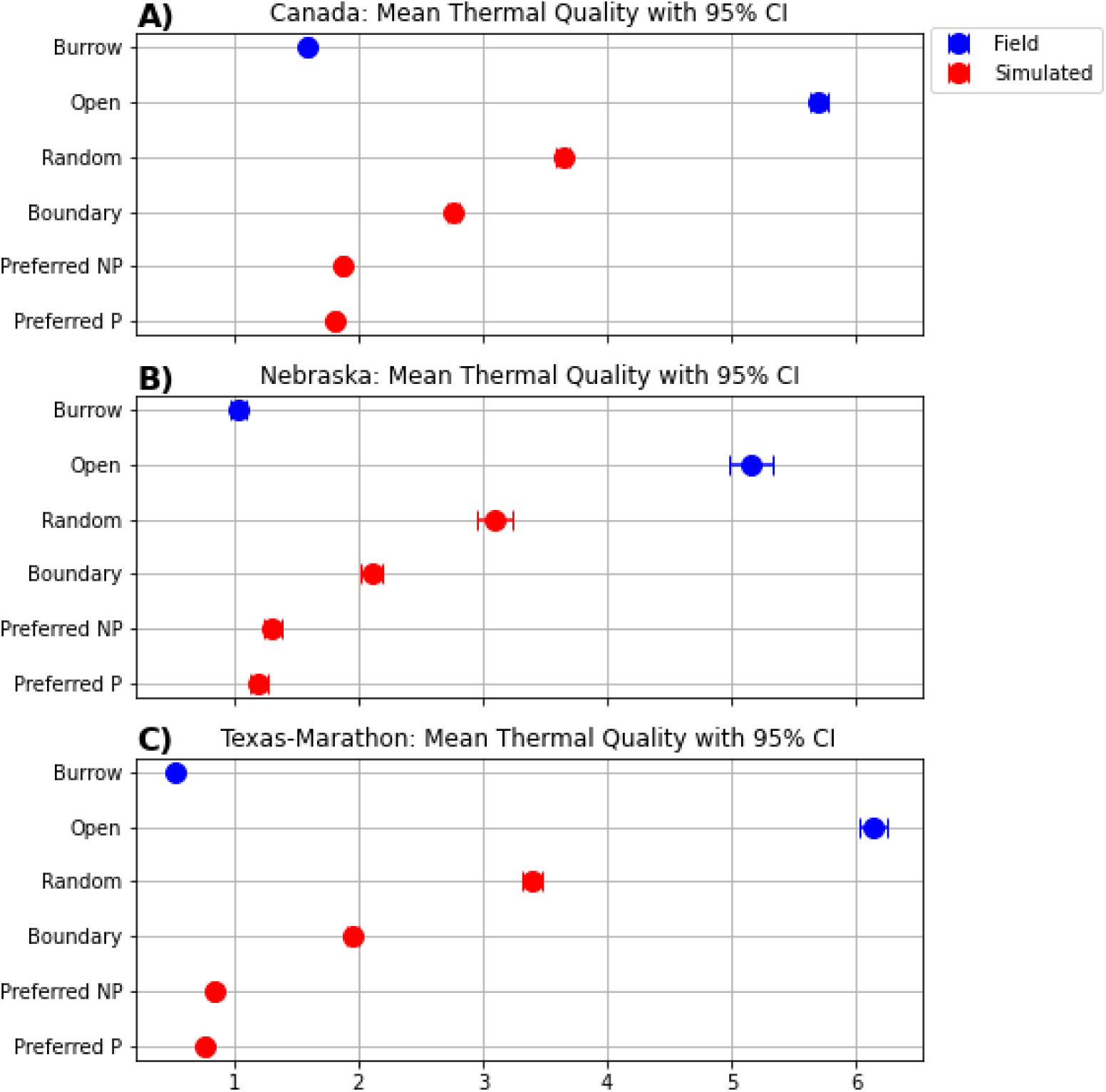
Thermal quality by microhabitat and for simulated snakes. Thermal quality (de) is a metric that measures the difference between environmental temperature (T*_e_*) from the thermal preference zone (Hertz et al. (1993), Piasečná et al. (2015), Taylor et al. (2021)). Thermal quality traditionally can not be estimated for snakes caught in the field due to not knowing what microhabitat (T*_e_*) they may be sampling. However, we can assess this for simulated agents to gain a better understanding of the thermal conditions snakes may experience in field conditions.

**Figure 9:**
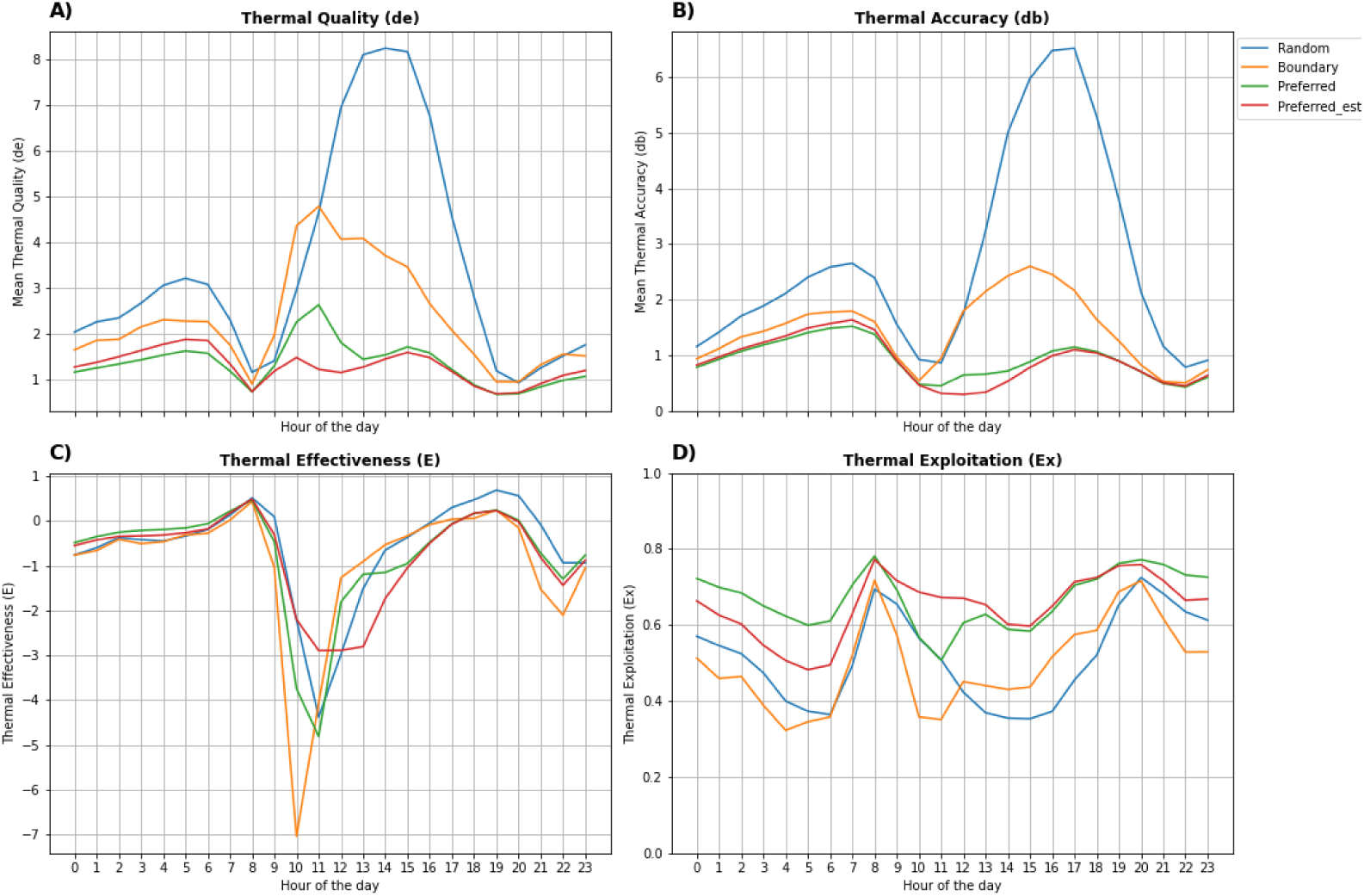
Plot grid of four thermal indices calculated by hour of the day. Figure A showcases thermal quality (de) is a metric that measures the difference of the thermal operative temperature (T*_e_*) from the thermal preference zone. So high values of de indicate low quality thermal habitat. Thermal accuracy (db) which is a metric that measures the difference between an organism’s body temperature (T*_b_*) from its thermal preference zone. High values of db indicate poor thermal accuracy. Figure C showcases thermal effectiveness (E) which is one minus the ratio of mean db and mean de. Values of one indicate careful and precise thermoregulation, zero indicate the animal does not need to thermoregulate and is thermoconforming, while values approaching negative one indicate periods of where the organism is in poor habitat and are actively avoiding thermoregulation. Finally, Figure D is the thermal exploitation index (Ex) which is the percentage of time spent in the thermal preference zone. Higher values indicate more time spent in the thermal preference zone (Hertz et al. (1993), Piasečná et al. (2015), Taylor et al. (2021))

**Figure 10:**
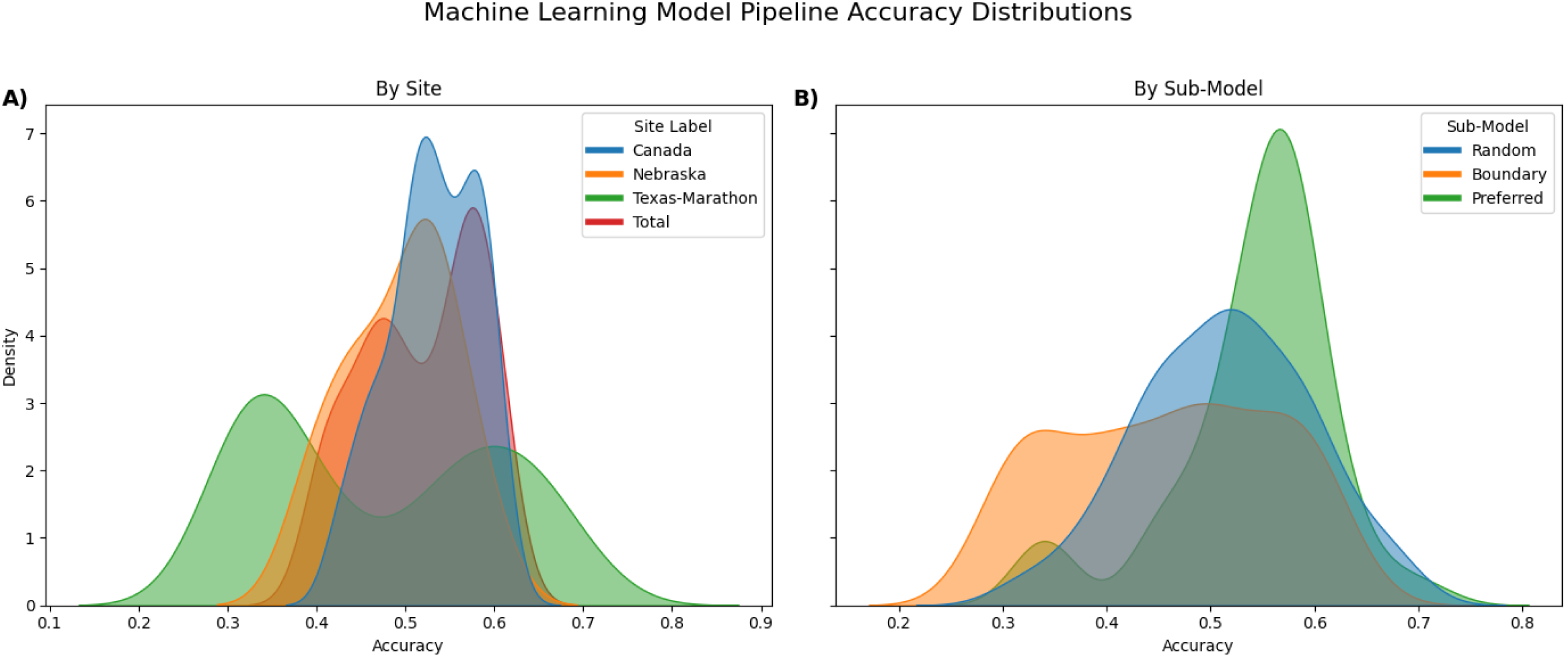
Kernel density plots of the accuracy results for all machine learning models built. Figure A is the results grouped by the sample population (Study Site) and Figure B is the results grouped by the thermoregulatory sub-model used to simulate training labels.

**Table 7:** The top 10 models from machine learning pipeline predicting microhabitat occupancy for all snakes sampled (study site = total population) under various model conditions. Filtering the data this way yields us models where we assume that intraspecific differences among populations can be generalized with a single model. The first five columns are the model specifications, columns 6-9 showcase model performance, and columns 10-14 are further details to how performance metrics were calculated. In our analysis we primarily focused on the accuracy of the model over metrics such as precision, recall, and F1 Score.

| Top 10 (Accuracy): Total Population Models |  |  |  |  |  |  |  |  |  |  |  |  |  |
| --- | --- | --- | --- | --- | --- | --- | --- | --- | --- | --- | --- | --- | --- |
| Sub-Model | Parameters Estimated? | Sampling Granularity (Min) | ML Method | Window Size | Accuracy | Precision | Recall | F1 Score | TP | FP | TN | FN | AUC |
| random | Yes | 15 | RF | NaN | 0.611020 | 0.541520 | 0.851103 | 0.661901 | 463 | 392 | 280 | 81 | 0.627439 |
| random | Yes | 70 | rnn | 9.0 | 0.610236 | 0.582278 | 0.410714 | 0.481675 | 46 | 33 | 109 | 66 | 0.610915 |
| preference | Yes | 70 | RF | NaN | 0.608365 | 0.560000 | 0.486957 | 0.520930 | 56 | 44 | 104 | 59 | 0.598913 |
| random | Yes | 70 | rnn | 4.0 | 0.606178 | 0.576471 | 0.426087 | 0.490000 | 49 | 36 | 108 | 66 | 0.627748 |
| preference | Yes | 15 | rnn | 7.0 | 0.605459 | 0.560360 | 0.571691 | 0.565969 | 311 | 244 | 421 | 233 | 0.617250 |
| random | No | 70 | rnn | 8.0 | 0.603922 | 0.588235 | 0.353982 | 0.441989 | 40 | 28 | 114 | 73 | 0.624704 |
| preference | Yes | 15 | rnn | 5.0 | 0.603633 | 0.553691 | 0.606618 | 0.578947 | 330 | 266 | 401 | 214 | 0.620327 |
| preference | Yes | 15 | rnn | 6.0 | 0.603306 | 0.554983 | 0.593750 | 0.573712 | 323 | 259 | 407 | 221 | 0.612152 |
| random | Yes | 70 | rnn | 10.0 | 0.600791 | 0.550000 | 0.495495 | 0.521327 | 55 | 45 | 97 | 56 | 0.653788 |
| random | No | 70 | rnn | 6.0 | 0.599222 | 0.556818 | 0.433628 | 0.487562 | 49 | 39 | 105 | 64 | 0.616581 |

## 4. Discussion

Our primary objective in this study was to examine the utility of a computational simulation framework for studying and quantifying the thermal effects of thermoregulatory behavioral strategies. Secondary goals were to evaluate potential trade-offs in thermal conditions associated with different algorithms, and explore the potential for our framework to classify patterns of microhabitat occupancy over time. From our analyses, both the boundary and preferred sub-models were relatively equivalent at reproducing real world conditions, while the preferred*_est_* consistently outperformed the other sub-models in time series similarity metrics (Table 3, 4, 5). This could indicate that individuals may adopt a mix of strategies depending on the environmental conditions, and thermoregulation strategy should not be treated as a static trait of the individual. Secondly, the superior performance of preferred*_est_* group indicates a potentially substantial role of individual variation when modeling thermal preferences.

Further evidence that thermoregulation strategies may be mixed can be qualitatively assessed by comparing simulated thermal accuracy to field snake values (Figure 7). At the Canada field site, both the preferred and boundary thermal accuracy values align with field conditions, with the preferred value corresponding to the lower thermal accuracy range and the boundary value matching snakes at the upper bounds. The preferred and the preferred*_est_* model thermal accuracy better matches field conditions compared to the boundary model at the Nebraska and Texas Site. One explanation for this pattern is that the Canada site had the poorest burrow thermal quality compared to the other two sites (Figure 8) which indicates individuals would need to spend more time thermoregulating in order to maintain preferred body temperatures. This would in turn lead to real conditions matching model assumptions more closely, as simulated snakes in our model only thermoregulate and do not hunt or rest, so a poorer average thermal quality would result in individuals prioritizing thermoregulation more than other behaviors potentially. From our thermal refugia analysis, the preferred sub-model has the fewest number of state flips (Figure 3) so from the perspective of an organism minimizing energetic costs, ectotherms in thermally favorable conditions would likely prefer the preferred strategy over the boundary strategy.

From a thermoregulatory perspective, organisms must maintain a balance between minimizing the energetic costs of switching microhabitat states and minimizing (db) under dynamic spatial-temporal temperature conditions. The main ecological trade-offs we identified between the boundary and preferred sub-models were during midday hours when thermal quality was at its worst regardless of strategy. Under these conditions, the boundary thermoregulatory submodel experienced the worst thermal quality (de); however, the boundary sub-model did not switch microhabitat states as often, which may minimize energy expenditure from a movement perspective in a living organism. All models had relatively equivalent thermal exploitation index values, even at midday. Even though the boundary thermoregulatory model resulted in the lowest thermal accuracy (Figure 9B). However, the thermal exploitation index being relatively equivalent across models (Figure 9C), the boundary strategy may be the most advantageous in the context of foraging as organisms following this strategy could spend more time exposed under thermally challenging conditions.

How individual different species or individual ectotherms navigate these trade-offs may partially explain why many ectotherms become cathermal under certain conditions, meaning environmental conditions primarily drive activity times of the organisms (rather than diel cycle) (Barbour and Clark (2012)). Results from our machine learning pipeline further corroborate this as our attempt to build supervised machine learning models to predict microhabitat occupancy from environmental variables proved largely unsuccessful (Figure 10). Our best model yielded an accuracy of 0.71 at one site, indicating that this modeling framework could have potential, but would likely require further integration of biotic and ecological phenomena such as predation risk, prey availability, or movement dynamics.

Fitness outcomes often hinge on trade-offs involving movement cost, energetics, and predation risk, none of which are currently included in our model. Although our model does allow us to identify the behavioral and temporal trade-offs of thermoregulatory patterns, it does not explicitly model fitness, which would be required for developing an understanding of how dynamics such as movement cost, energetics, spatial thermal heterogeneity, and other risks play into thermoregulatory decisions. This is also likely the explanation for why our sub-models produce relatively accurate body temperature values, but we had inconclusive results for predicting microhabitat occupancy as occupancy may be shaped primarily by selection pressures and a variety of environmental conditions rather than solely temperature and thermoregulation.

## 5. Conclusion and Future Work

The notion that organisms adopt strategies to navigate the fundamental fitness trade-off faced by ectotherms—namely balancing the physiological cost of poor thermal accuracy and the energetic costs of thermoregulation (e.g., seeking thermal refuges) is a long-established idea (Huey and Slatkin (1976), Vickers et al. (2011)). Our framework aims to find application with this theory to provide a methodology that elucidates how, when, and why ectotherms choose various behavioral thermoregulation strategies.

Our research highlights the potential of a simulation framework for understanding more nuanced thermoregulatory behavior, but also indicates that simulations of thermoregulatory behavior or microhabitat use may be more accurate if they can incorporate more behavioral and ecological complexity. Future iterations of our model could explicitly model fitness as a function of not only thermal accuracy, but also incorporate factors such as movement cost, hunger, and risk to analyze how individuals navigate these trade-offs and quantify the emergent patterns at the population level.

Our simulation modeling approach and overarching modeling pipeline (Figure 1) provide researchers with a tool to analyze these conditions at both fine temporal resolutions and site-specific scales—without necessarily requiring the collection of live organisms. Such an analysis would be invaluable for predicting how organisms may adapt to climate change, as plasticity in thermoregulatory behavior will almost certainly mediate how ectotherm survival and reproduction are affected by shifting temperatures (Buckley et al. (2015)). In conjunction, our method could be leveraged as a pre-hoc OTM analysis leveraging just microhabitat data to inform field site selection and assess whether functional differences in thermoregulation are likely among different populations of ectotherms.

Lastly, future work could expand on the potential of simulation frameworks for understanding individual variation in thermal ecology. The best performing sub-model (preferred*_e_ st*) assumed individuals differed in thermal preference, suggesting that there is significant intraspecific variation in body temperature dynamics of the individual snakes in our study. Intraspecific variation in thermal traits has also been linked to personality traits in poikilotherms, suggesting that modeling certain thermoregulatory characteristics (e.g., low thermal accuracy) is correlated with bolder (high-risk taking strategies) behavioral syndromes in lizards (Horváth et al. (2024)). The boundary sub-model as the bold strategy, as it results in higher exposure rates at the cost of poorer thermal accuracy, implying that a promising direction for future work is to make thermoregulatory behavior selection a function of “safe” (the preferred sub-model) versus “risky” (the boundary sub-model) agent strategies.

